# On-farm biomass recycling with biostimulant Re-Gen increases corn yields in multi-year farm trials

**DOI:** 10.1101/2024.04.12.589288

**Authors:** William S Gibson, Amy S Ziobron, Noah E Olson, Deborah A Neher, Charles F Smith, Victoria I Holden

## Abstract

The United States produced 15.1 billion bushels of corn for grain in 2023, relying on harmful synthetic chemicals such as fertilizer and pesticides to ensure high yields. This dependency on agrochemicals has negatively impacted the environment and soil microbiome, therefore, there is a need to rebuild soil health by implementing regenerative agricultural practices. One increasingly utilized regenerative practice is the application of biostimulants, or microbial inoculants that can rebuild soil health and productivity. A multi-year corn trial was conducted to quantify the impact of Re-Gen, a biostimulant invented to degrade woody biomass and increase nutrient bioavailability in the soil, to increase corn yield at a dairy farm in Ferrisburgh, Vermont. Over the two-year trial, Re-Gen application on corn stover and cover crop residues increased corn bushels per acre by 24% and increased tons of corn silage per acre by 12.5-30%, depending on the field. Soil nutrient analysis and plant tissue analysis showed increased nutrients, particularly in one trial field. Multi-year Re-Gen application showed increased monetary value, indicating that the effects of Re-Gen do not diminish with multiple applications. Further investigation into the mechanism suggests that increased phosphatase production stimulated by Re-Gen could contribute to increased phosphorus bioavailability in the soil and uptake in the tissue, potentially increasing yields. These results highlight the potential for Re-Gen to foster regenerative agriculture processes while also increasing yield and, therefore, revenue for corn farmers in the United States.

## 1 Introduction

More than 15.1 billion bushels of corn for grain were produced in the United States in 2023 alone (USDA, 2023). To grow such large amounts of corn, the use of synthetic chemical inputs is often required, including fertilizers and pesticides. In Midwestern corn and soybean cropping systems, where more than 75% of all US corn is grown, nitrogen fertilizer is applied at an average rate of 135 kg N/ha/year, amounting to $15.8B spent on fertilizer in 2022 in the Midwestern US alone (Russell et al., 2009;USDA, 2023;USDA and US Agricultural Statistics Service, 2023). The dependency upon agrochemicals, and other synthetic inputs, has ultimately negatively impacted the environment and soil microbiome of the Midwest (Russell et al., 2009;Revillini et al., 2019;Potter et al., 2022) and increased financial strain on farmers due to rising costs and demand for nutrients (USDA, 2022). Soil productivity is particularly a concern for agricultural regions in the United States responsible for the production of high demand commodity crops, namely corn and soybean (Hatfield et al., 2013). As farmers wish to reduce reliance on harmful agricultural chemicals in their growing practices, there is a need to rebuild soil health while protecting farmer health and maintaining crop yields.

Regenerative agricultural practices aim to mitigate farmer dependence on synthetic inputs and conventional pesticides, while increasing resilience and sustainability of the global food production system (Muhie, 2022). Many regenerative and organic practices further aim to restore natural processes, bridge gaps in nutrient cycling, and promote ecosystem services (McLennon et al., 2021). Thermophilic composting is one practice that utilizes natural microbial decomposition for the recapture of nutrients from managed organic wastes (Grand, 2020). While compost improves soil health and serves as a rich amendment of microorganisms and nutrients, this strategy also requires upfront resource, time, and labor investments, which may not be feasible for farmers at certain production scales (Wright et al., 2022;Olson et al., 2024). Even for those making these upfront investments, the consistency and succession of microbial populations from composting systems can still vary (Neher et al., 2013;Olson et al., 2024). Additional regenerative practices for improving soil health and reducing nutrient loss are cover cropping and leaving corn residues in-field after harvest, called corn stover (Adetunji et al., 2020). Cover crops and corn stover mitigate erosion and improve soil biological characteristics through increasing organic matter and available nutrients, further stimulating microbial abundance and diversity (Melkonian et al., 2017;Adetunji et al., 2020;Nalley and Lee, 2023). However, like composting, cover crop and stover systems require proper management to ensure farmers receive soil health benefits. Without proper management, cover crop residues and corn stover can sequester or tie up nutrients, decreasing or reducing nutrient availability for the subsequent crop (Melkonian et al., 2017;Adetunji et al., 2020). Ultimately, regenerative solutions such as composting and cover cropping can increase soil heath and improve sustainability within food production systems (Adetunji et al., 2020;Khangura et al., 2023). However, access to resources, time, or labor investments may remain as barriers for the adoption of such practices at large agricultural scales, or for those who do not need additional synthetic inputs.

Being mindful of farmer reliance upon agrochemicals, despite an increased demand for sustainable food production, there is a need for environmentally friendly solutions that increase farm profitability and can be adopted easily at all growing scales and alongside conventional inputs. An emerging class of non-chemical products aimed at addressing this need is biostimulants (Albrecht, 2019). Biostimulants are substances or microorganisms applied to plants and/or soils for purposes ranging from facilitation of nutrient uptake efficiency, protection against pathogens, increasing stress tolerance, or improving characteristics of crop quality (du Jardin, 2015). Microbial inoculants, either as a single microorganism or collection of species, are a common form of plant biostimulants currently available on the market. Re-Gen is one such commercially available microbial inoculant composed of a consortium of naturally occurring, beneficial soil microorganisms. The inoculant was developed by IMIO Technologies, Inc. (Essex Junction, VT) to accelerate the in-field degradation of lignin-rich and woody biomass, mainly through Re-Gen’s three primary functional groups: 1) Lignin degrading microbes, to degrade tough plant fibers, increasing the nutrient potential of residual biomass (Kumar and Chandra, 2020); 2) Lactic acid producers, to increase the rate of biodegradation and improve soil organic matter content (Raman et al., 2022); and 3) Plant Growth Promoting Rhizobacteria (PGPR), to develop symbiotic relationship between plants and microbes and facilitate increased nutrient uptake and protection against pathogens (de Andrade et al., 2023). The capacity for Re-Gen to degrade lignin-rich plant material into a high nutritive value growing substrate was evaluated previously in *Cannabis sativa*, which showed an increase in the production of oxidase and phenol oxidase, oxidative enzymes facilitating lignin degradation, in the presence of Re-Gen (Olson et al., 2024). Additionally, nutrient analyses of the microbial treated hemp biomass revealed profiles similar to those of finished compost (Olson et al., 2024). Together these findings suggest Re-Gen-treated plant matter may serve as an organic fertilizer substitute, based on the inoculant’s ability to access the nutrient potential of tough fibers through ligninolytic enzyme production.

Based on the findings demonstrated in *Cannabis sativa*, the ability for IMIO Re-Gen to increase bioavailable nutrients for subsequent crops was further evaluated on corn. In collaboration with an independent agronomist and 500-cattle Vermont dairy farm, Re-Gen was applied in-field to stover and cover crop residues prior to planting feed corn, at multiple field sites in Ferrisburgh, VT over two years. It was hypothesized that the field-application of Re-Gen could increase corn yield through accelerated degradation of cover crop residuals; specifically, by increasing the pool of bioavailable nutrients in the soil for the subsequent corn crop. The impacts of Re-Gen at the field scale were evaluated according to soil and plant tissue analyses, as well as total yield expressed as silage weight.

## 2 Materials and Methods

### 2.1 Farm and Trial Information

Nea Tocht Farm is an 800-acre, 500-head dairy cattle farm in Ferrisburgh, VT. The property consists of land formations and soils typical of Addison County on the West side of US Route 7. Most of the soils are heavy Vergennes or Covington clays, dotted with “mounds” of stony silt-loam, such as Melrose, or sandy-loam such as Adams. The farm family owners-operators keep themselves apprised of current information and technology, and farm with the care of keeping the herd and land as healthy and resilient as possible over time. They use no-till, conservation tillage, and strategic full tillage, as well as cover cropping and crop rotation strategies. Liquid dairy manure is the main source of applied nutrients. Several types of organic residuals such as wood ash, and paper fiber as well as calcitic lime are used in conjunction with the farm’s NRCS 590-compliant Nutrient Management Plan, as required within the Vermont Required Agricultural Practices. This farm family has been exploring how to enhance soil health, soil functions, nutrient cycling efficiencies, and reducing the use of chemical inputs such as herbicides, seed treatments, and conventional fertilizers.

Year 1 of a two-year corn trial spanned May to October 2022. The four fields used in Year 1 of the trial were VA01, BH01, VA04, and LA05. For each field, 0.5 acres total were included in the trial. The biostimulant Re-Gen was applied on 0.25 acre for each field directly to the in-field cover crop residues on May 18, 2022. The remaining 0.25 acre of control fields were left untreated. Seedway 3750 corn was planted on 5/25/2022 and 5/27/2022 at a seeding rate of 34,000 seeds per acre, and the crop was managed with minimal interventions throughout the season. Corn was harvested between 6-8 October 2022, and sample data for each treatment and field was collected by harvesting one contiguous row representing 1/1000^th^ of an acre. Weight per plant (ear weight for one ear per plant) was recorded to calculate total plant weight, total ear weight, and bushels per acre. Once weighed, the 1/1000th acre ear samples were chopped, and collected samples for each treatment group were sent to an accredited agricultural feed laboratory (Dairy One, Inc., Ithaca, NY) for forage feed analysis, as feed for commercial dairy cattle. Wet ear weight was converted to bushels per acre using the Method for Hand Harvested Ear Corn described by the University of Wisconsin Agronomy Department (Wright et al., 2022).

To better address our trial partner’s needs, our 2023 trial focused on corn silage yield. Year 2 of the two-year corn trial spanned May to October 2023. Two fields used in Year 1 of the trial, VA04 and LA05, were treated in Year 2 at a total of 1 acre. The biostimulant Re-Gen was applied on 0.5 acre total for each field, and each field was split into two equal 0.25 acre treatment groups: “Re-Gen: 1 year,” in which Re-Gen was applied to cover crop residues for the first time; and “Re-Gen: 2 year,” in which Re-Gen was applied to cover crop residues for the second consecutive year. The biostimulant application date was 22 May 2023. The remaining 0.25 acre of control fields were left untreated. Seedway 3750 corn was planted on 8 June 2023 at a seeding rate of 34,000 seeds per acre, and the corn was managed using minimal interventions throughout the season. Corn was harvested for silage from 2 to 7 October 2023 and sample data for each treatment and field was collected by harvesting three separate contiguous rows, each representing 1/1000^th^ of an acre. Weight of whole plants in the 1/1000^th^ acre samples was recorded to calculate total silage per acre. Once weighed, the 1/1000th acre ear samples were chopped, and collected samples for each treatment group were sent to an accredited agricultural feed laboratory (Dairy One, Inc.) for commercial dairy cattle forage feed analysis, including milk pounds per ton of dry matter. This figure is often used to determine corn silage feed quality (Shaver, 2007). To determine the overall milk pounds that could be generated by feeding cows with corn silage harvested from control- or Re-Gen-treated fields, the milk pounds per ton of dry matter was multiplied by the tons of corn silage determined from the yield data.

Growing season statistics for both years of the trial are detailed in Table 2 (CLIMOD;DeGaetano et al., 2016). In 2023, Vermont faced unprecedented flooding in July (25.4 cm or 10.02 inches) that negatively impacted farm yields throughout the state, most commonly damaging crops meant for feed (Banacos, 2023;Watson, 2023). Because the soils are predominantly clay-based, most of the Nea Tocht Farm fields were super-saturated with some ponding for much of the month of August and into September. The Re-Gen trial plots and control areas were not ponded at any time.

**Table 1:**
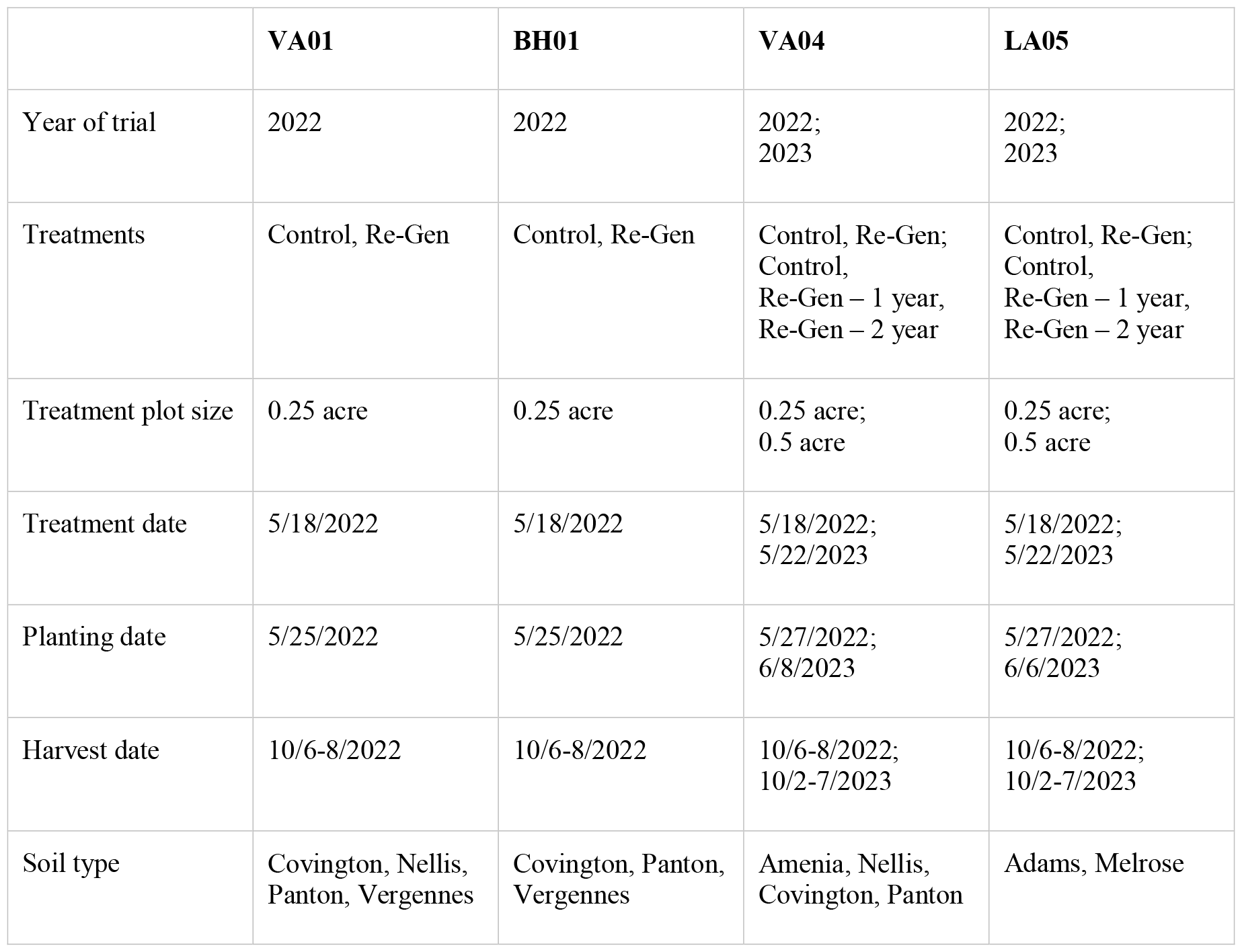
Experimental information for the Re-Gen farm trial at Nea Tocht Farm in Ferrisburgh, VT.

**Table 2:**
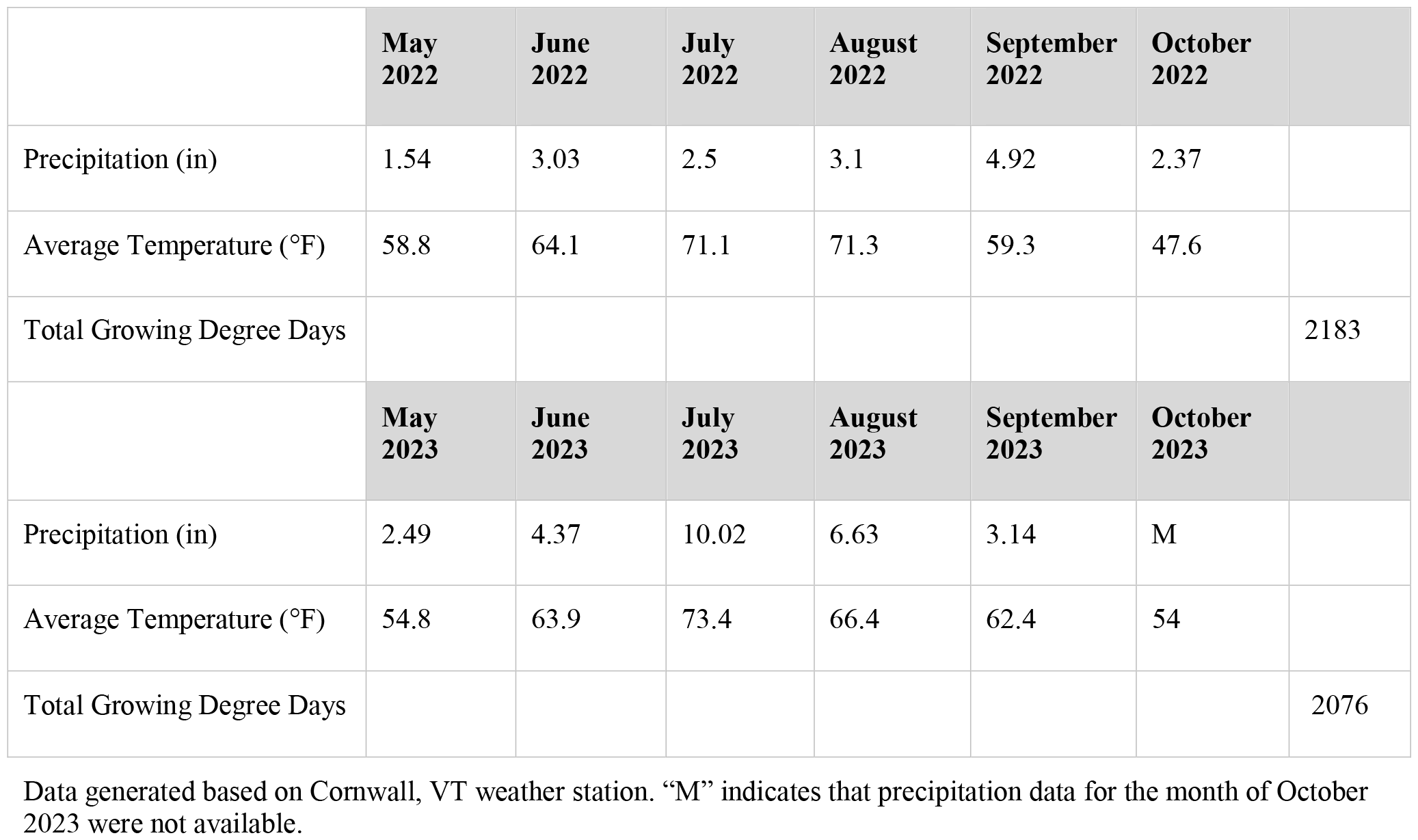
Seasonal weather data collected in 2022 and 2023.

### 2.2 Biostimulant Preparation

Re-Gen (IMIO Technologies, Inc.) was prepared according to the manufacturer’s directions. In Year 1, liquid Re-Gen was activated with provided microbial nutrient media for 4-8 hours, then further diluted to 30-35 gallons of water/acre prior to in-field application. In Year 2, approximately 12g of Re-Gen powder per acre was reanimated in 8 ounces of water per acre and allowed to sit for 16-24 hours to activate the product. Reanimated Re-Gen was then further diluted into 30-35 gallons of water per acre for application.

### 2.3 Soil analysis

At the conclusion of the 2023 growing season, soil samples were collected from each treatment field using the Zig-Zag method as described by the University of Vermont Extension Agricultural and Environmental Testing Lab (UVM, 2022). Samples were sent for soil analysis by the Maine Soil Testing Service. Nutrients were extracted using a Modified Morgan nutrient extraction procedure, allowing extraction of all major nutrients and micronutrients (McIntosh, 1969).

### 2.4 Plant Tissue analysis

In the tassel to silking stage, samples of plant tissue were collected by choosing representative leaves at ear nodes for each treatment condition in the 2023 trial. Samples were sent for plant tissue analysis to Agro One / Dairy One using Service Package 180 that tests total nitrogen, phosphorus, potassium, calcium, magnesium, sulfur, zinc, copper, iron, boron, and manganese (DairyOne, 2023).

### 2.5 Phosphatase activity

In previous experiments conducted and described by Olson, et al 2024, leftover organic hemp biomass was treated with 1 mL of Re-Gen per gram of organic matter and incubated for 21 days. Control samples were treated with 1 mL of water per gram of organic matter and incubated for 21 days. Samples were collected at day 7, 14, and 21 to determine phosphatase activity using previously published protocol in which 0.3 □g of shredded hemp was mixed in 200 □ mL of autoclaved nanopure water (Neher et al., 2017;Olson et al., 2024).

## 3 Results

### 3.1 2022 and 2023 Trial Results

In 2022, control-treated fields averaged 138.7 bushels per acre while Re-Gen treated fields averaged 171.9 bushels per acre, an increase of 24% or 33 bushels (Figure 1).

**Figure 1:**
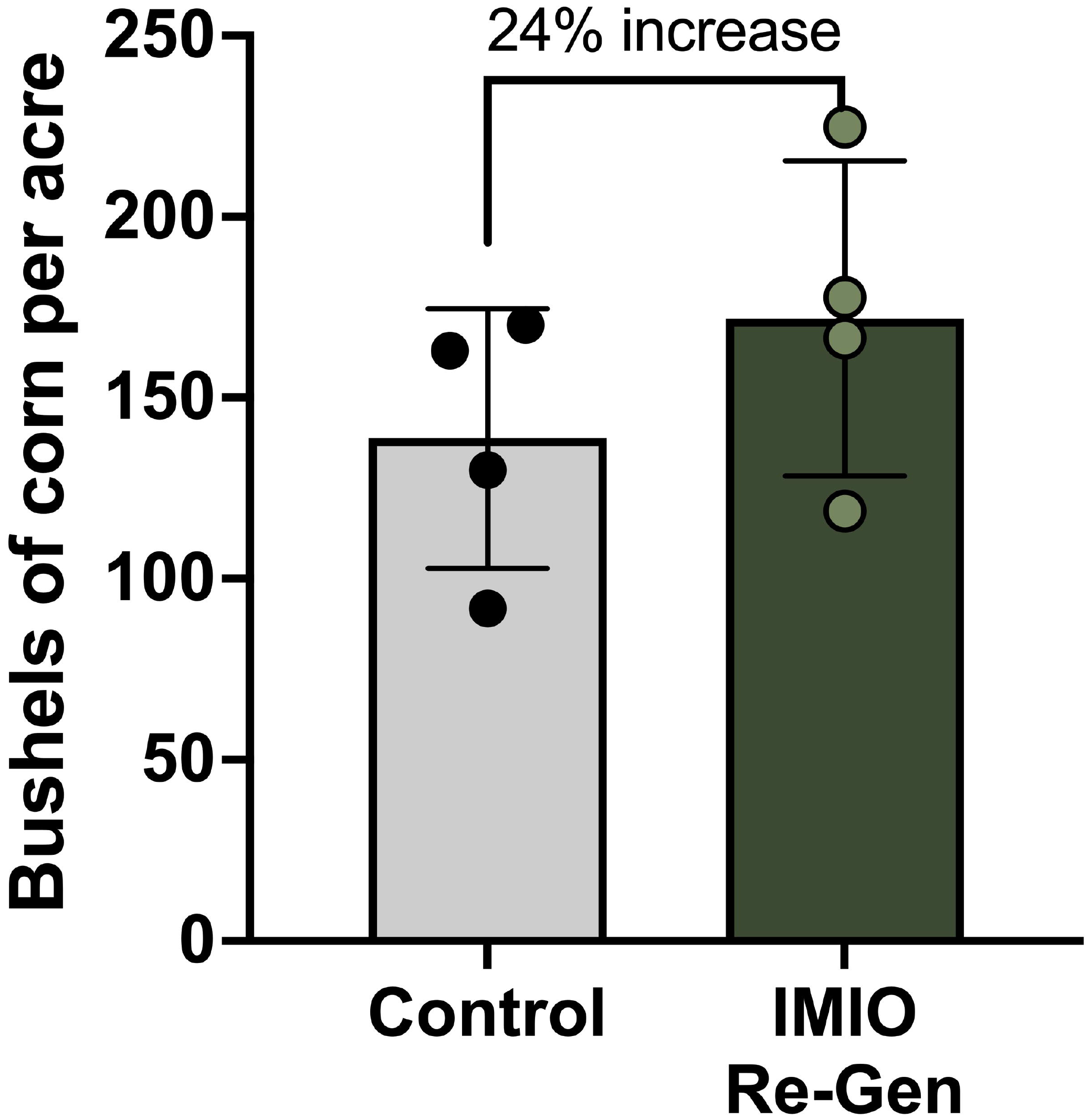
Fields treated with Re-Gen saw a 24% increase in bushels per acre compared to control-treated fields. VA01, BH01, VA04, and LA05 were treated with control or Re-Gen 2 weeks prior to planting corn. Upon harvesting, plant and ear mass was collected for each treatment in each field. Wet ear mass was converted to bushels using the Wisconsin method for Hand Harvesting Ear Corn. Each data point represents one biological replicate corresponding to a distinct field.

In 2023, control treatment in fields LA05 and VA04 resulted in 14.14 and 12.48 tons/acre, respectively (Figure 2). First-time treatment with Re-Gen (“Re-Gen: 1 year”) resulted in 18.29 and 14.03 tons/acre, an increase of 30% and 12.5%, respectively, compared to control fields. Second-time treatment with Re-Gen (“Re-Gen: 2 year”) resulted in 16.89 and 15.76 tons/acre, an increase of 19% and 17%, respectively, compared to control fields.

**Figure 2:**
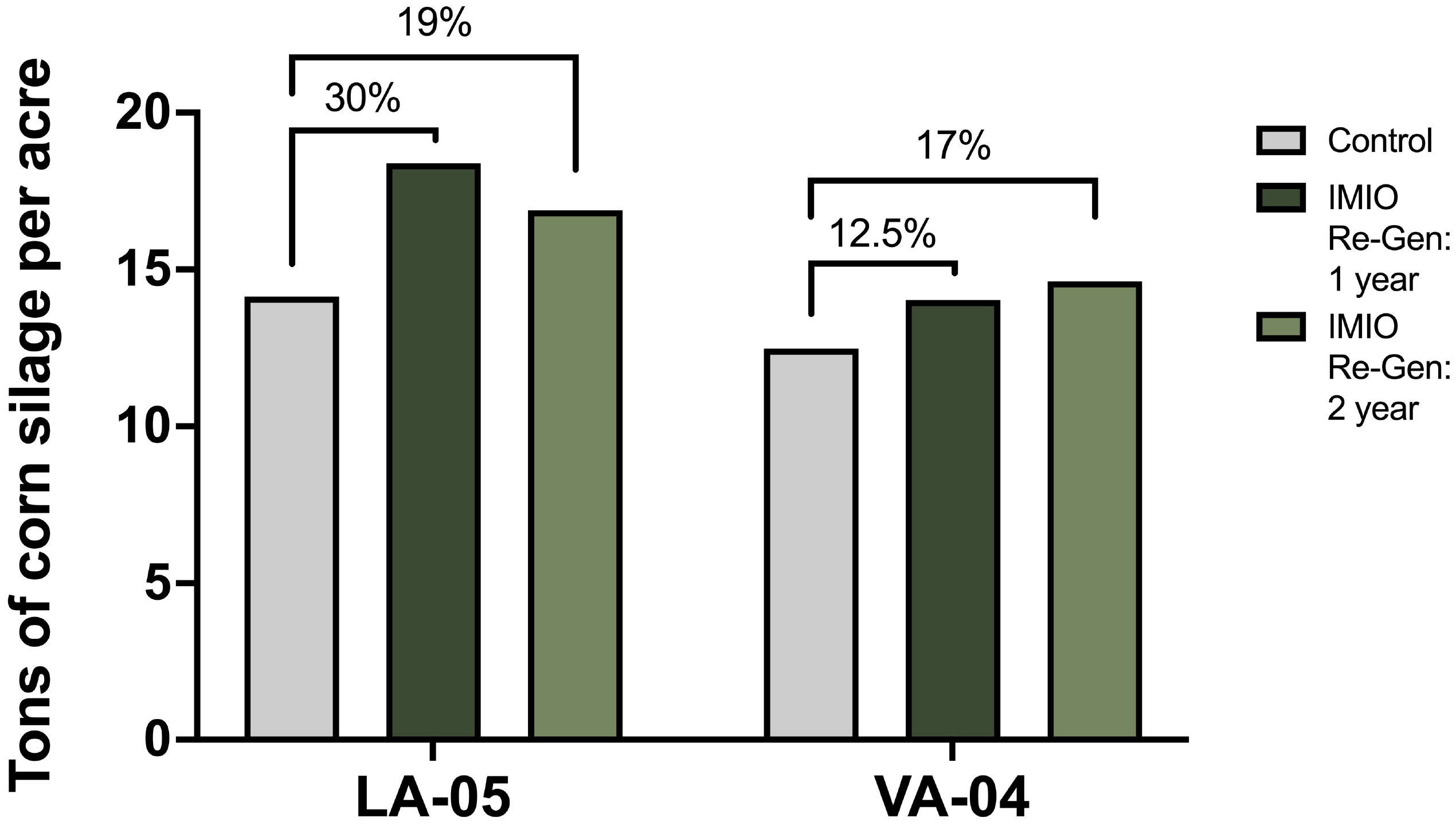
Fields treated with Re-Gen saw an increase in corn silage compared to control fields, an increase seen in consecutive treatment years. VA04 and LA05 were treated with control or Re-Gen 2 weeks prior to planting corn. Upon harvesting, corn silage was collected for each treatment at 3 sites in each field. Each bar represents the average of the three sites for each treatment.

Soil nutrient analysis results, including optimal nutrient levels for corn as provided by Maine Soil Testing Service, are shown in Table 3. Most nutrients are within the optimal levels for corn for each field, however, there are substantial differences between LA05 and VA04 nutrient concentrations, such as in pH, organic matter %, and most notably phosphorus (as phosphate in lb/acre).

**Table 3:**
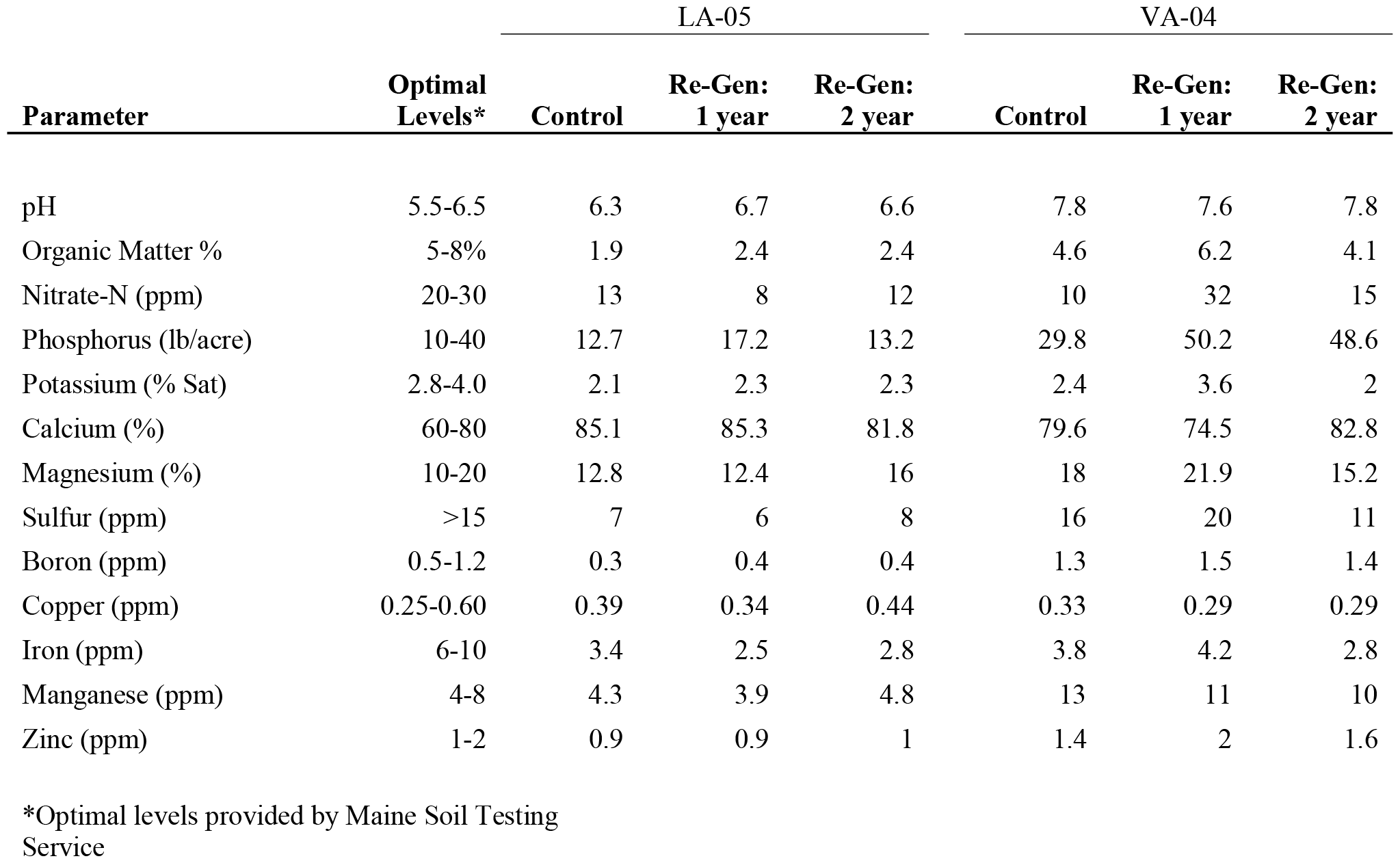
Properties of soil following 1 or 2 years of Re-Gen treatment.

Plant tissue nutrient results and sufficiency levels for silking as provided by Cornell University Cooperative Extension are shown in Table 4. Most plant nutrients are within the sufficiency levels for corn for each field, however, there are noteworthy nutrients that were deficient or nearly deficient in the plant tissue for control-treated samples from field VA04 that were increased upon 1- or 2-year Re-Gen treatment: nitrogen, phosphorus, copper, and zinc (Table 4). These results were only seen in field VA04, highlighting the differences in the two fields within the 2023 trial (Table 1).

**Table 4:**
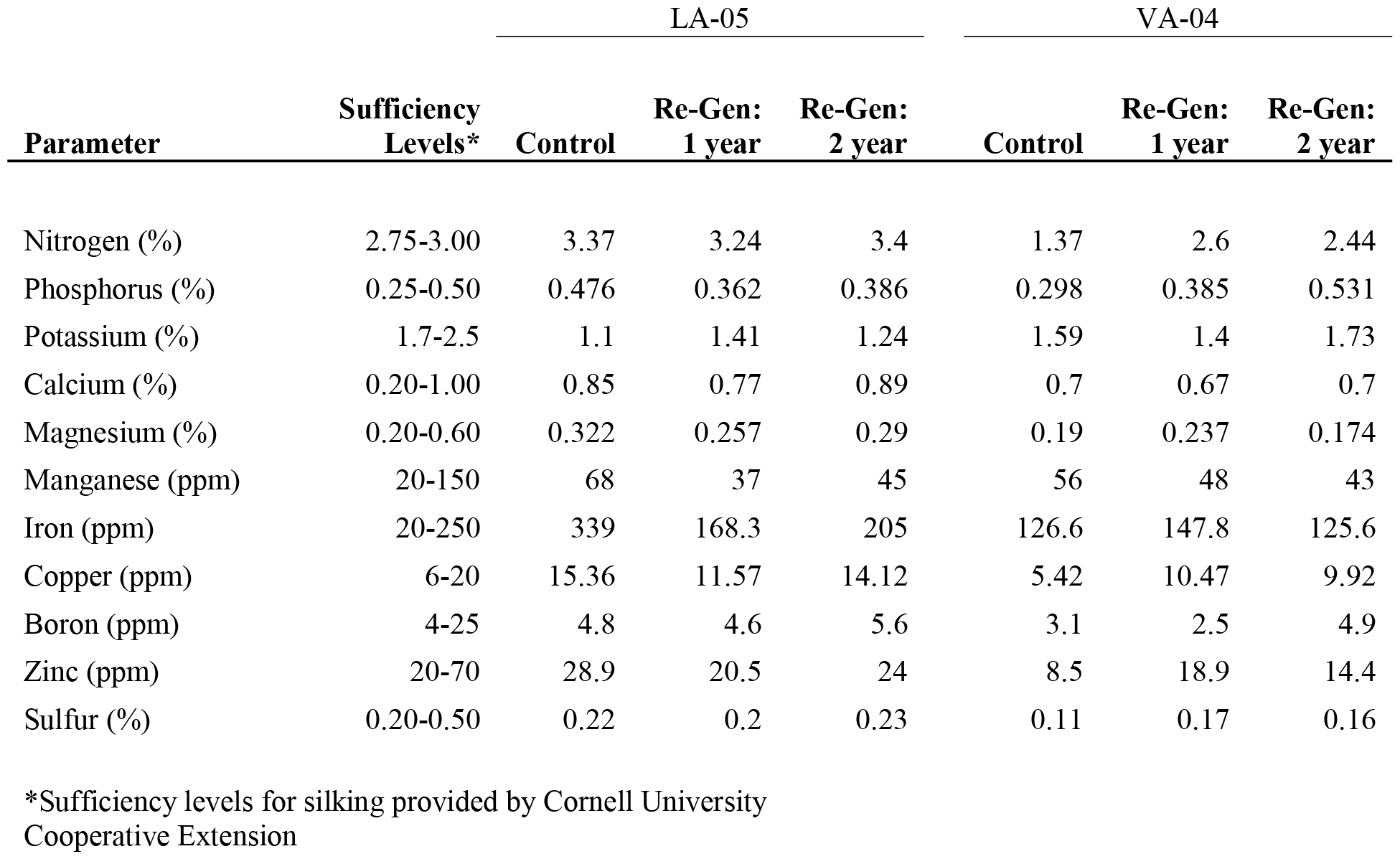
Leaf tissue analysis results following control and Re-Gen treatment.

### 3.2 Phosphorus Bioavailability

Sufficient phosphorus is important for high yield crops, but an excess of phosphorus in the soil can lead to this nutrient moving off-site and causing water quality issues. As noted above, treatment of cover crop residues with Re-Gen for 1 or 2 years increased phosphorus bioavailability in the soil (Figure 3A) and in plant tissue (Figure 3B), particularly for field VA04. Re-Gen treated biomass had elevated phosphatase activity at days 6, 14, and 21 compared to the control that was treated with water, indicating that the microbes within Re-Gen are able to produce phosphatases (Figure 4). These phosphatases could be partially responsible for the phosphorus results described in Tables 2 and 3 and highlighted in Figure 3.

**Figure 3:**
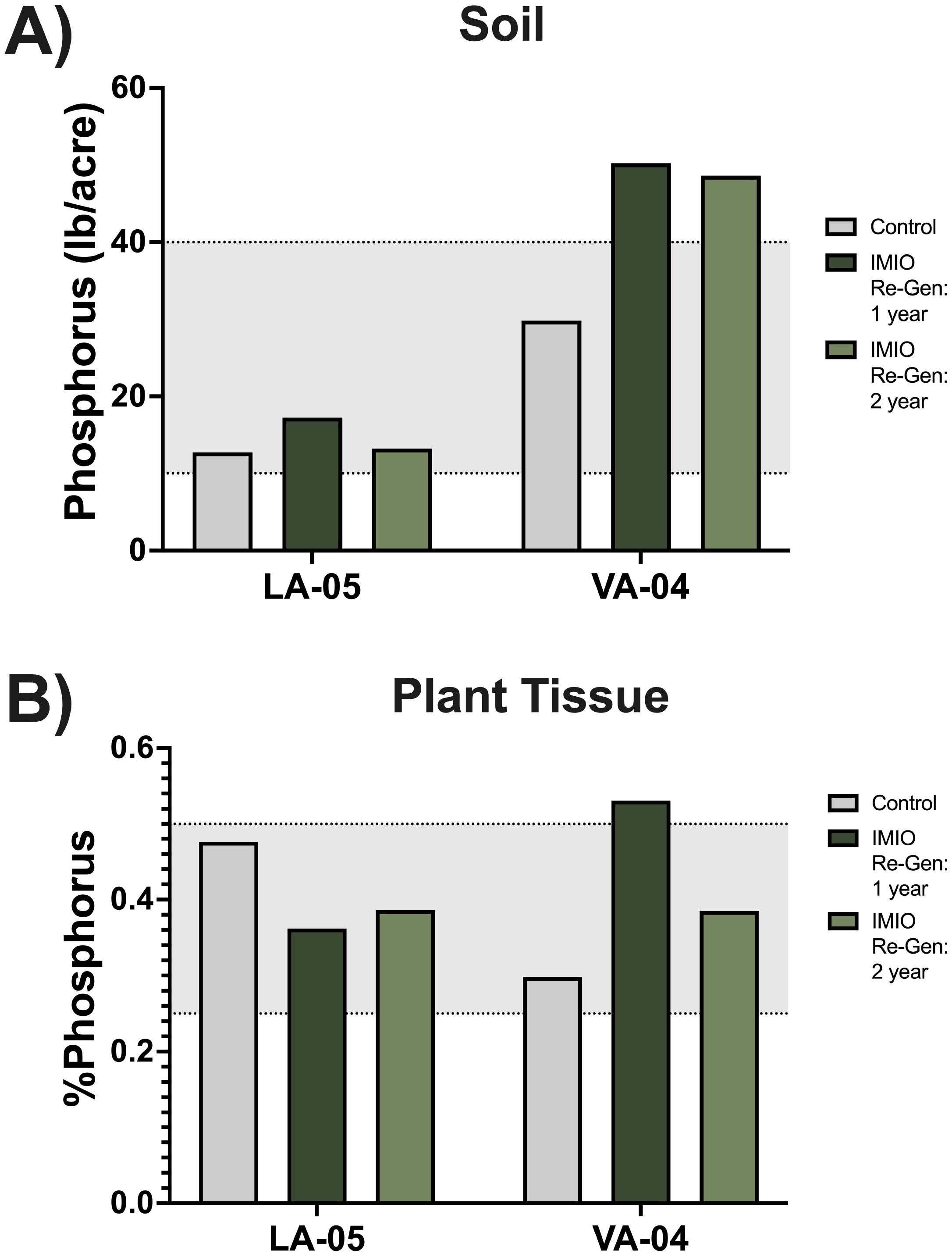
Phosphorus in soil and plant tissue is elevated after Re-Gen treatment in field VA-04. End of season soil samples (A) and silking plant tissue samples (B) were collected from soil and plants in fields LA05 and VA04. Each bar represents one sample. Gray shading represents optimal and sufficiency levels for soil and plant tissue, respectively.

**Figure 4:**
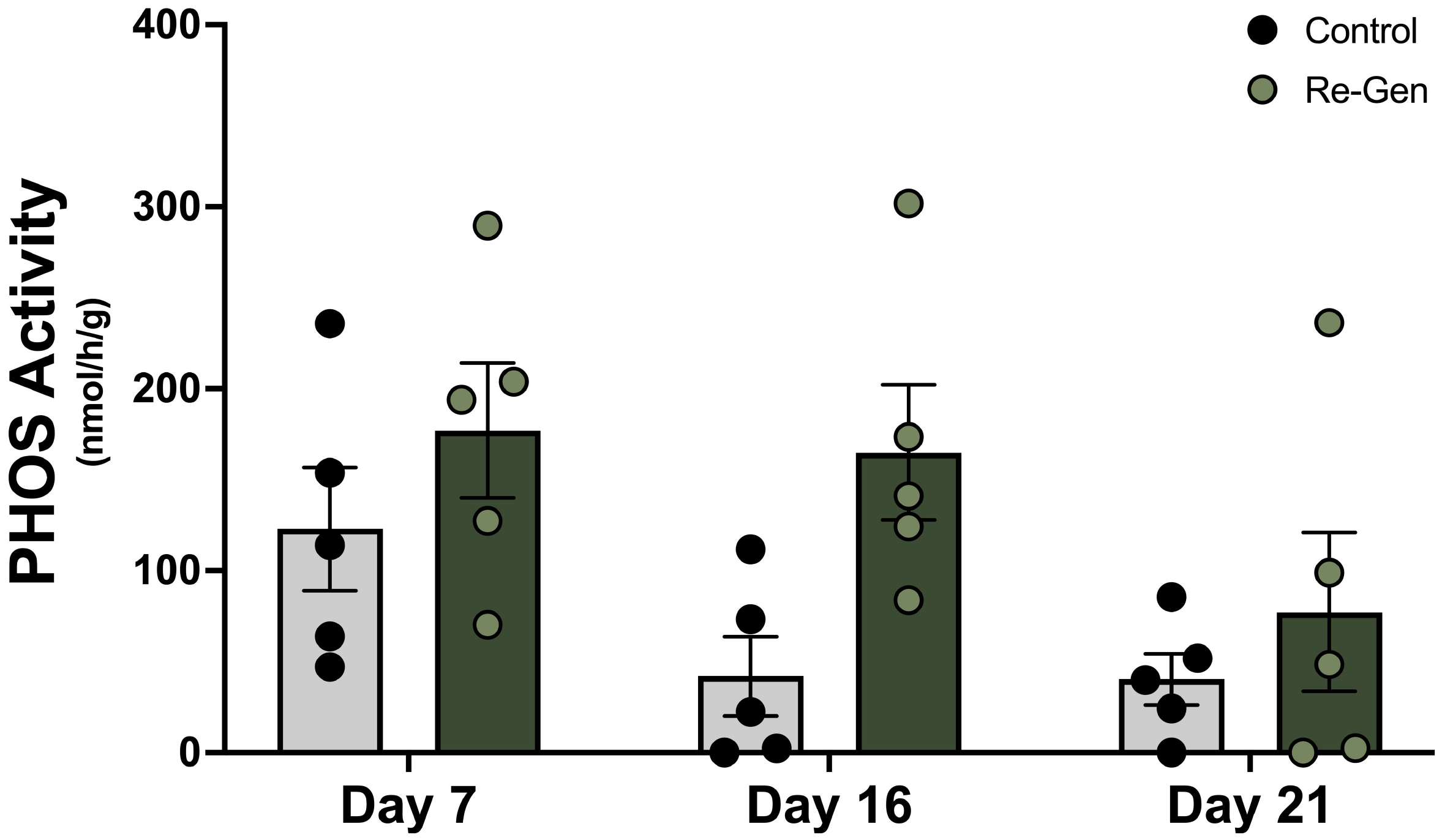
Leftover organic biomass treated with Re-Gen showed more phosphatase activity compared to control-treated biomass. Phosphatase enzyme activity was measured at days 7, 16, and 21 on organic hemp biomass that was recycled with water (control) or Re-Gen. Each data point represents one technical replicate.

### 3.3 Economic Impact of Re-Gen Application

In our 2022 trials, there was a 24% increase in bushels per acre, equaling a 33-bushel increase. Considering that the 2022 average price of a bushel of corn was $6.54, this represents a $215.82 increase per acre. Considering the cost of Re-Gen is $18 per acre, this is a 12-fold return on investment.

These data indicate a potential 24% or 20% increase in total milk pounds for silage harvested from field LA05 with 1- or 2-year Re-Gen treatment, respectively, compared to control (Figure 5A). There was also a more moderate increase in total milk pounds for silage harvested from field VA04 of 9% or 7% upon 1- or 2-year Re-Gen treatment compared to control (Figure 5B). Together, these data indicate the potential for a considerable economic gain when Re-Gen is used to recycle cover crop residues in fields prior to planting corn.

**Figure 5:**
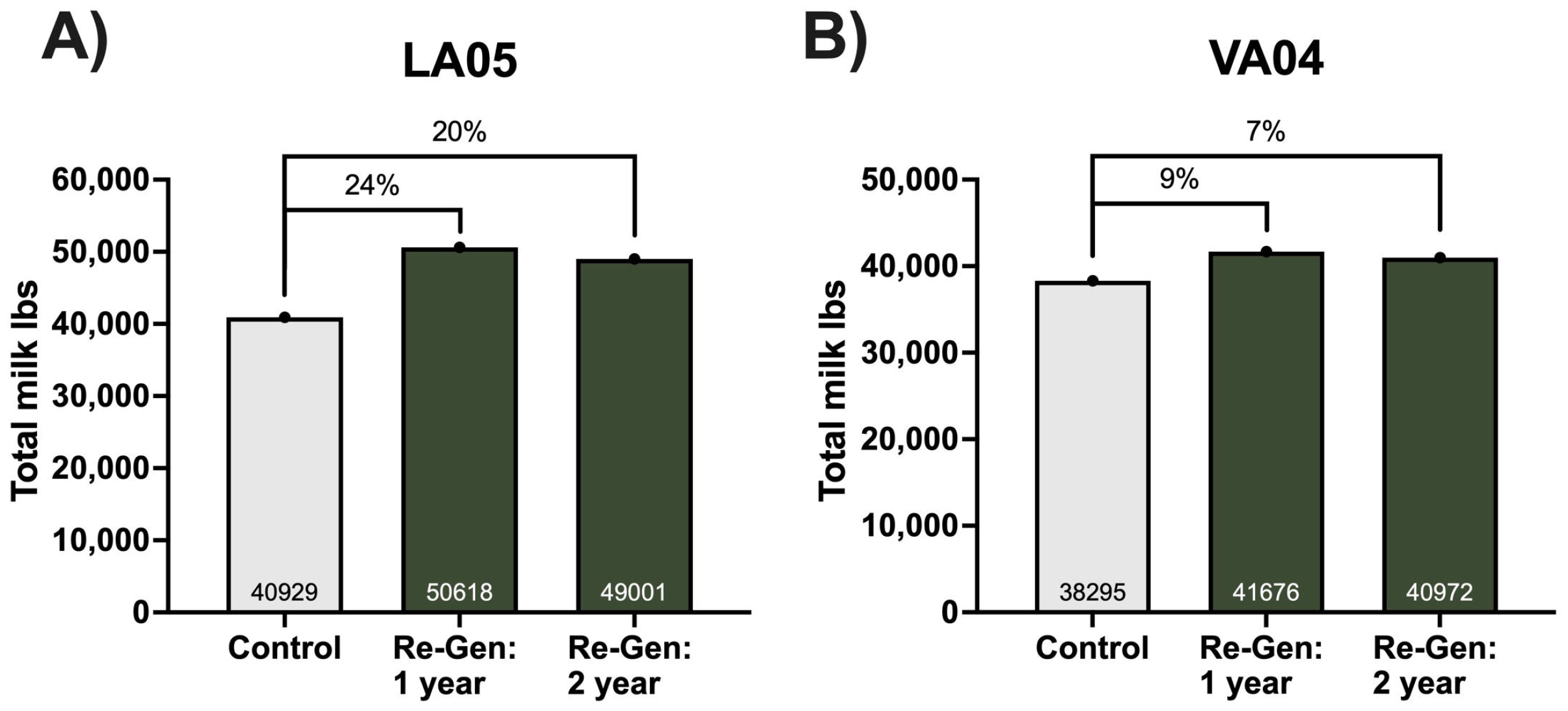
Treatment with Re-Gen can increase the economic value of corn silage as reported by total milk pounds. Total milk pounds for 2023 corn silage harvest were calculated by multiplying the number of milk pounds per ton, determined by Dairy One corn silage analysis, by the yields in tons per acre.

## 4 Discussion

Our study supported our hypothesis that in-field residue treatment with Re-Gen would increase corn yield. Yield increases were directly related to increased nutrient bioavailability, as evidenced by soil nutrient analysis as well as plant tissue analysis data. The increased yield may be influenced by increased phosphorus availability, a hypothesis that is supported by previous studies showing elevated phosphatase activity in Re-Gen-treated organic biomass. These effects translate into greater net income for farmers, indicated by an increase in revenue of bushels per acre in 2022 as well as increased total milk production in 2023.

One pattern evident in the data is the variability in results between the two fields used in the 2023 trial, LA05 and VA04. Although these two fields are both part of the larger Nea Tocht Farm, they are located in different areas and have contrasting soil characteristics. LA05 is a higher elevation field with more sandy soil, whereas the soil in VA04 is variable throughout, but generally is a heavier soil with more clay. Even though these fields have unique characteristics, both fields saw double-digit increases in yield upon Re-Gen treatment when compared to control. However, the pattern of yield increases was different: LA05’s yield increase was larger for 1^st^ year treatment of Re-Gen compared to the 2-year treatment, whereas VA04 saw a larger yield increase for 2^nd^ year Re-Gen treatment compared to the 1^st^ year treatment. These data could indicate that richer, higher-yielding fields like LA05 could have a large spike in yield from Re-Gen treatment during the first year of Re-Gen implementation that lessens or plateaus with multi-year treatment, whereas less rich, lower-yielding fields may have compounding benefits the longer that Re-Gen is used on the field. Additional research is necessary to better understand the benefits of Re-Gen treatment on cover crop residues on different types of fields and over extended multi-year treatments.

Re-Gen treated organic biomass produces more phosphatase enzyme compared to control biomass, which correlates positively with elevated phosphorus in the soil in the 2023 farm trial. Phosphorus has the positive impact of supporting corn growth but can have a negative environmental impact if applied in excess. Phosphorus is essential for nearly all stages of corn growth, from root development and early growth to reproductive development of the corn plant (Pereira et al., 2020). Because of its importance, phosphorus is commonly applied at planting to ensure adequate yields at a rate of 89.7 – 134.5 kg per hectare (80-120 pounds per acre), depending on geographic location and soil characteristics (LaBarge, 2022). However, excessive application of phosphorus results in dangerous runoff into streams and lakes that results in eutrophication (Conley et al., 2009). Another factor contributing to phosphorus runoff is the presence of non-labile phosphorus, or phosphorus that is unavailable for plant uptake due to a variety of factors including immobilization, adsorption, and soil pH and/or mineralogy. Non-labile phosphorus requires processes like desorption and mineralization of organic matter by soil microbes, including by phosphatase processing, for phosphorus to become available for uptake (Prasad and Chakraborty, 2019). Many states, including Vermont, Ohio, and Minnesota, have instituted programs to reduce phosphorus runoff into bodies of water, such as Lake Champlain and the Great Lakes (H2Ohio, 2024;MDA, 2024;NRCS, 2024). If Re-Gen application can increase phosphorus bioavailability in the soil, this could result in additional phosphorus uptake (immobilization) by the plant, potentially preventing phosphorus runoff into nearby lakes and streams, or a decreased need for phosphorus to be applied in-field. More work needs to be done to characterize the contribution of Re-Gen recycled biomass and its potential impact on phosphorus uptake and runoff.

While these data are encouraging for Vermont dairy and corn farmers, further trials need to be performed to understand the impact of Re-Gen on farms in the Midwestern United States, where the vast majority of corn farming takes place. These trials, currently underway, will investigate the effectiveness of Re-Gen in the region as well as the feasibility of incorporating Re-Gen into current farming protocols. One concern that will be addressed in these studies is the ability of Re-Gen to be mixed with common herbicides, such as glyphosate (Round-Up). In the Midwestern United States, herbicide burndown is a common cover crop termination method, used to mitigate nutrient competition and ensure a release of any scavenged nutrients back to the field (Jewett and Thelen, 2007;Palhano et al., 2018;Adetunji et al., 2020). A widely used commercial herbicide for this strategy is glyphosate (Kumar et al., 2023). While herbicidal desiccation of residual cover crop biomass increases soil nutrient levels (Jewett and Thelen, 2007), the use of glyphosate has also been associated with decreased soil biological activity and microbial diversity (Busse et al., 2001;Nguyen et al., 2016), as well as adverse non-target effects on human and environmental health (Kanissery et al., 2019). Therefore, if a farmer is able to mix Re-Gen with glyphosate, it could help restore the microbial population while increasing nutrient bioavailability following cover crop burndown. Furthermore, the logistical benefit of mixing Re-Gen with glyphosate is that it requires one pass through the fields rather than two, saving labor and fuel costs. This method of application would keep costs low while restoring the microbial ecosystem and generating additional revenue from their corn harvest.

Overall, this study illustrates the potential yield gains made possible by applying biostimulants to cover crop residues, resulting in increased nutrient recycling and bioavailability in corn production. Additionally, this study highlights the potential for biostimulants to increase sustainability within conventional cropping systems while increasing economic value by mitigating the need for excess synthetic nutrients and increasing soil microbial life. More work must be done to incorporate regenerative practices into agriculture to create a more sustainable agricultural ecosystem.

## 5 Conflict of Interest

Amy Ziobron is an employee of IMIO Technologies, Inc. and holds equity in the company. Charles Smith and Victoria I. Holden are co-founders of IMIO Technologies, Inc. and hold partial ownership in the company.

## 6 Author Contributions

WG designed the experimental setup, performed field trials, collected and analyzed data, and contributed to manuscript composition. AZ contributed to manuscript composition. CS contributed to manuscript revisions. VH analyzed data and contributed to manuscript composition.

## 7 Funding

Re-Gen development and characterization was funded by NSF SBIR Grant 2014792. The analyses in this study were paid for by IMIO Technologies, Inc.

## 8 Acknowledgments

We would like to thank the Vander Wey Family of Nea Tocht Farm for trialing Re-Gen on their fields, applying the product, and sharing their observations during the 2022 and 2023 farm trials.

## 9 Data Availability Statement

The original contributions presented in the study are included in the article; further inquiries can be directed to the corresponding author.

## References

Adetunji, A., Ncube, B., Mulidzi, R., and Lewu, F. (2020). Management impact and benefit of cover crops on soil quality: A review. Soil and Tillage Research 204.

Albrecht, U. (2019). “Plant Biostimulants: Definition and Overview of Categories and Effects”. (UF/IFAS Extension: University of Florida).

Banacos, P. (2023). The Great Vermont Flood of 10-11 July 2023: Preliminary Meteorological Summary [Online]. National Weather Service: National Oceanic and Atmospheric Administration. Available: https://www.weather.gov/btv/The-Great-Vermont-Flood-of-10-11-July-2023-Preliminary-Meteorological-Summary [Accessed 2024].

Busse, M., Ratcliff, A., Shestak, C., and Powers, R. (2001). Glyphosate toxicity and the effects of long-term vegetation control on soil microbial communities. Soil Biology and Biochemistry 33, 1777–1789.

Climod Northeast RCC CLIMOD 2 [Online]. ACIS NOAA Regional Climate Centers. Available:http://climod2.nrcc.cornell.edu/ [Accessed].

Conley, D.J., Paerl, H.W., Howarth, R.W., Boesch, D.F., Seitzinger, S.P., Havens, K.E., Lancelot, C., and Likens, G.E. (2009). Controlling eutrophication: nitrogen and phosphorus. Science 323, 1014–1015.

Dairyone (2023). Plant Tissue Analysis [Online]. Available:https://dairyone.com/services/forage-laboratory-services/plant-tissue-analysis/ [Accessed].

De Andrade, L.A., Santos, C.H.B., Frezarin, E.T., Sales, L.R., and Rigobelo, E.C. (2023). Plant Growth-Promoting Rhizobacteria for Sustainable Agricultural Production. Microorganisms 11.

Degaetano, A., Moore, R., Belcher, B., and Eck, B. (2016). CSF Growing Degree Day Calculator [Online]. Climate Smart Farming: Cornell University. Available:http://climatesmartfarming.org/tools/csf-growing-degree-day-calculator/ [Accessed 2024].

Du Jardin, P. (2015). Plant biostimulants: Definition, concept, main categories and regulation. Scientia Horticulturae 196, 3–14.

Grand, A. 2020. Compost: Thermophilic Compost. Available:https://www.best4soil.eu/assets/factsheets/6.pdf.

H2ohio (2024). Enhancing Water Quality and Improving Public Health [Online]. Available:https://h2.ohio.gov/ohio-epa/ [Accessed 2023].

Hatfield, J., Cruse, R., and Tomer, M. (2013). Convergence of agricultural intensification and climate change in the Midwestern United States: implications for soil and water conservation. Mar Freshw Res 64, 423–435.

Jewett, M., and Thelen, K. (2007). Winter cereal cover crop removal strategy affects spring soil nitrate levels. Journal of Sustainable Agriculture 29.

Kanissery, R., Gairhe, B., Kadyampakeni, D., Batuman, O., and Alferez, F. (2019). Glyphosate: Its Environmental Persistence and Impact on Crop Health and Nutrition. Plants (Basel) 8.

Khangura, R., Ferris, D., Wagg, C., and Bowyer, J. (2023). Regenerative Agriculture - A Literature Review on the Practices and Mechanisms Used to Improve Soil Health. Sustainability 15.

Kumar, A., and Chandra, R. (2020). Ligninolytic enzymes and its mechanisms for degradation of lignocellulosic waste in environment. Heliyon 6, e03170.

Kumar, V., Singh, V., Flessner, M.L., Haymaker, J., Reiter, M.S., and Mirsky, S.B. (2023). Cover crop termination options and application of remote sensing for evaluating termination efficiency. PLoS One 18, e0284529.

Labarge, G. (2022). Developing Phosphorus and Potassium Recommendations for Field Crops. Agriculture and Natural Resources.

Mcintosh, J. (1969). Bray and Morgan Soil Extractants Modified for Testing Acid Soils from Different Parent Materials1. Agronomy Journal 61, 259–265.

Mclennon, E., Dari, B., Jha, G., Sihi, D., and Kankarla, V. (2021). Regenerative agriculture and integrative permaculture for sustainable and technology driven global food production and security. Agronomy Journal 113.

Mda (2024). Minnesota Agricultural Water Quality Certification Program [Online]. Minnesota Department of Agriculture. Available:https://www.mda.state.mn.us/environment-sustainability/minnesota-agricultural-water-quality-certification-program [Accessed 2024].

Melkonian, J., Poffenbarger, H., Mirsky, S., Ryan, M., and Moebius-Clune, B. (2017). Estimating Nitrogen Mineralization from Cover Crop Mixtures using the Precision Nitrogen Management Model. Agronomy Journal 109, 1944–1959.

Muhie, S. (2022). Novel approaches and practices to sustainable agriculture. Journal of Agriculture and Food Research 10.

Nalley, R., and Lee, C. (Year). “Investigating the Nitrogen Penalty of Cereal Cover Crops in No-till Corn (Zea mays L.)”, in: ASA CSSA SSSA International Annual Meeting).

Neher, D., Fang, L., and Weicht, T.R. (2017). Ecoenzymes as indicators of compost to suppress Rhizoctonia solani. Compost Science & Utilization 25, 251–261.

Neher, D.A., Weicht, T.R., Bates, S.T., Leff, J.W., and Fierer, N. (2013). Changes in bacterial and fungal communities across compost recipes, preparation methods, and composting times. PLoS One 8, e79512.

Nguyen, D., Rose, M., Rose, T., Morris, S., and Van Zwieten, L. (2016). Impact of glyphosate on soil microbial biomass and respiration: A meta-analysis. Soil Biology and Biochemistry 92, 50–57.

Nrcs (2024). Conservation Stewardship Program [Online]. Natural Resources Conservation Service. Available:https://www.nrcs.usda.gov/programs-initiatives/csp-conservation-stewardship-program [Accessed].

Olson, N., Neher, D., and Holden, V. (2024). On-Farm Conversion of Cannabis sativa Waste Biomass into an Organic Fertilizer by Microbial Digestion. Compost Science & Utilization.

Palhano, M., Norsworthy, J., and Barber, T. (2018). Evaluation of Chemical Termination Options for Cover Crops. Weed Technology 32, 227–235.

Pereira, N., Galindo, F., Gazola, R., Dupas, E., Rosa, P., Mortinho, E., and Filho, M. (2020). Corn Yield and Phosphorus Use Efficiency Response to Phosphorus Rates Associated With Plant Growth Promoting Bacteria. Frontiers in Environmental Science 8.

Potter, T.S., Vereecke, L., Lankau, R.A., Sanford, G.R., Silva, E.M., and Ruark, M.D. (2022). Long-term management drives divergence in soil microbial biomass, richness, and composition among upper Midwest, USA cropping systems. Agriculture, Ecosystems & Environment 325.

Prasad, R., and Chakraborty, D. (2019). “Phosphorus Basics: Understanding Phosphorus Forms and Their Cycling in Soil”, (ed.) A.a.a.U. Extension.).

Raman, J., Kim, J.S., Choi, K.R., Eun, H., Yang, D., Ko, Y.J., and Kim, S.J. (2022). Application of Lactic Acid Bacteria (LAB) in Sustainable Agriculture: Advantages and Limitations. Int J Mol Sci 23.

Revillini, D., Wilson, G.W.T., Miller, R.M., Lancione, R., and Johnson, N.C. (2019). Plant Diversity and Fertilizer Management Shape the Belowground Microbiome of Native Grass Bioenergy Feedstocks. Front Plant Sci 10, 1018.

Russell, A.E., Cambardella, C.A., Laird, D.A., Jaynes, D.B., and Meek, D.W. (2009). Nitrogen fertilizer effects on soil carbon balances in midwestern U.S. agricultural systems. Ecol Appl 19, 1102–1113.

Shaver, R. (2007). “Evaluating Corn Silage Quality for Dairy Cattle”. University of Wisconsin Extension).

Usda (2022). “Impacts and Repercussions of Price Increases on the Global Fertilizer Market”, in: International Agricultural Trade Report.).

Usda (2023). “Crop Production”. (Washington, D.C.: Crop Reporting Board, Statistical Reporting Service).

Usda, and Us Agricultural Statistics Service (2023). “Farm Production Expenditures: 2022 Summary”. (Washington, D.C.: USDA).

Uvm (2022). How to Take a Soil Sample [Online]. University of Vermont Extension. Available:https://www.uvm.edu/sites/default/files/UVM-Extension-Cultivating-Healthy-Communities/how-to-take-a-soil-sample-2.pdf [Accessed 2024].

Watson, D. (2023). “Severe Weather and Flooding Loss & Damage Survey Results”. Vermont Agency of Agriculture, Food & Markets).

Wright, J., Kenner, S., and Lingwall, B. (2022). Utilization of Compost as a Soil Amendment to Increase Soil Health and to Improve Crop Yields. Open Journal of Soil Science 12.

